# Simulating lesion-dependent functional recovery mechanisms

**DOI:** 10.1101/2021.01.20.427450

**Authors:** Noor Sajid, Emma Holmes, Thomas M. Hope, Zafeirios Fountas, Cathy J. Price, Karl J. Friston

**Affiliations:** Wellcome Centre for Human Neuroimaging, University College London, 12 Queen Square, London, UK; Emotech Labs, London, UK

**Keywords:** functional recovery, active inference, degeneracy, functional reorganisation, experience-dependent plasticity

## Abstract

Functional recovery after brain damage varies widely and depends on many factors, including lesion site and extent. When a neuronal system is damaged, recovery may occur by engaging residual (e.g., perilesional) components. When damage is extensive, recovery depends on the availability of other intact neural structures that can reproduce the same functional output (i.e., degeneracy). A system’s response to damage may occur rapidly, require learning or both. Here, we simulate functional recovery from four different types of lesions, using a generative model of word repetition that comprised a default premorbid system and a less used alternative system. The synthetic lesions (i) completely disengaged the premorbid system, leaving the alternative system intact, (ii) partially damaged both premorbid and alternative systems, and (iii) limited the experience-dependent plasticity of both. The results, across 1000 trials, demonstrate that (i) a complete disconnection of the premorbid system naturally invoked the engagement of the other, (ii) incomplete damage to both systems had a much more devastating long-term effect on model performance and (iii) the effect of reducing learning capacity within each system. These findings contribute to formal frameworks for interpreting the effect of different types of lesions.

## 1. Introduction

Most patients who suffer functional impairments after brain damage improve over time. This has been demonstrated in motor (Connolly *et al.*, 1994; Langhorne *et al.*, 2009; Bultmann *et al.*, 2014), visual (Seghier *et al.*, 2005; Guzzetta *et al.*, 2010) and language (Saur *et al.*, 2006; Hope *et al.*, 2019). The degree of functional recovery is highly variable and lesion-dependent (Irle, 1987; Chen *et al.*, 2000; Alstott *et al.*, 2009). Here, we distinguish between two distinct lesion-dependent recovery mechanisms. For a premorbid system with partial damage, recovery may entail re-learning that increases the functional capacity of the damaged system—and may involve peri-lesional activity (Warburton *et al.*, 1999). In a severely damaged system, recovery depends on whether an alternative system can be used to reproduce the same functional output (Welbourne *et al.*, 2011; Seghier *et al.*, 2012).

The ability to use alternative systems for the same task is referred to as degeneracy (Tononi *et al.*, 1999; Price and Friston, 2002). For example, when reading written words aloud, with regular spelling to sound relationships, sounds can be generated either by using learnt spelling-to-sound relationships or by whole word recognition. If the predominant word recognition system fails, readers can instead use spelling-to-sound relationships. At a neurological level, degeneracy supports functional recovery, following damage to components of the premorbid system, by enabling intact cortical regions and pathways to recapitulate the same function. The new system may engage distinct components, relative to the premorbid system, but may share undamaged components (Seghier *et al.*, 2012). Like the reading example, different components may not have the same function in isolation, but together can produce the same output as the premorbid system. Degenerate systems may therefore support better behavioural performance than relying on a partially damaged premorbid system. However, when the lesion is extensive or affects multiple areas; all available degenerate systems may also be rendered dysfunctional.

If two neural systems for the same function are equally efficient and sufficient, then switching from one system to another could occur instantaneously. Conversely, if one system is preferred over another then it takes time to attain premorbid levels of proficiency — and may require functional reorganisation mediated by experience-dependent plasticity: i.e., re-learning due to behavioural experience (Nudo, 2003; Kleim and Jones, 2008; Fu and Zuo, 2011; Lövdén *et al.*, 2013; Nudo, 2013). During re-learning, experience-dependent plasticity helps to restore and compensate for functional deficit (Cooper, 2005). These learning processes depend on multiple mechanisms manifest at different temporal scales: namely, short-term plasticity mechanisms that occur rapidly (e.g., neuromodulatory changes in synaptic efficacy) and slower long-term reorganisation that support functional recovery (e.g., long-term potentiation and depression) (Kleim and Jones, 2008). These might be complemented by changes in the neural structure (Hope *et al.*, 2017) and/or electrophysiology (Carmichael, 2003). Previous models of plasticity-related recovery have shown re-learning can retune and realise the contribution of peri-lesional regions (Ueno *et al.*, 2011; Welbourne *et al.*, 2011).

Using *in-silico* lesions in a computational model of word repetition (i.e., repeating heard words), we characterise two functional recovery mechanisms. The first is the rapid recruitment of an alternative system that can reproduce the same outcome (in the context of degeneracy); the second is long-term plasticity within either the premorbid or alternative system. The recovery mechanism triggered is expected to depend on the structural, and implicitly computational, resources available to perform the task. We modelled these processes using active inference, which treats perception and action as belief updating under a particular generative model of the environment. Central to this approach is the notion that behaviour is Bayes optimal under some prior beliefs (following the complete class theorem). This allows us to characterise patients with brain damage as operating under (possibly lesional) priors that constitute their generative model; priors are statistical contingencies encoded by model parameters (e.g., synaptic connection strengths). Additionally, active inference provides a formal way of measuring: synaptic connectivity changes during learning, the accompanying neurophysiology (Friston *et al.*, 2017a) and degeneracy. Here, degeneracy is the entropy of Bayesian beliefs about the causes of sensations and redundancy is a measure of how much beliefs need to change to explain current observations (Sajid *et al.*, 2020c).

Building on prior work (Sajid *et al.*, 2020b; Sajid *et al.*, 2020c), we used a simulated work repetition paradigm to measure degeneracy, redundancy, and task performance under four different levels of *in-silico* lesion severity. The key extension in our current model of word repetition, compared to our previous model, is that we introduced two distinct degenerate networks—premorbid and alternative—into a hierarchical model that also captured the neuromodulatory aspect of attention. This allowed us to simulate the effects of different types of lesions that were not investigated in our prior work. The premorbid network represents the preferred (i.e., predominant) circuitry, prior to lesioning(Hickok, 2014; Hope *et al.*, 2014). The alternative network is less experienced but has the capacity to learn.

The resulting hierarchical model was lesioned in four ways: First, we simulated changes in model performance when the premorbid system was completely disconnected, and output depended on the less experienced alternative system. Second, we repeated Lesion 1 but added a second lesion to the intrinsic connectivity of the alternative system. This limited recovery by reducing the capacity of the alternative system to learn, over time, via experience-dependent plasticity. Third, rather than completely disconnecting the premorbid system, as in Lesions 1 and 2, we partially disconnected both the premorbid and alternative systems. This made it difficult for the higher level to identify which context (premorbid or alternative) would be most effective. Finally, we repeated Lesion 3 (partial disconnections to both systems) with an additional lesion to reduce relearning capacity.

Below, we briefly introduce active inference, generative models and the types of in-silico lesions that can be induced. In the following sections, we describe the word repetition model, how we use it to simulate functional recovery mechanisms and the results. Finally, to make quantitative and disambiguating predictions for future empirical work, we simulated electrophysiological responses that have already been associated with a decrease in baseline neuronal firing (Schiene *et al.*, 1996; Carmichael, 2003), and attenuated post-synaptic sensitivity (Luhmann *et al.*, 1995; Neumann-Haefelin *et al.*, 1995).

## 2. Active Inference

Active inference postulates that the brain self-organises by optimising two complementary objectives: 1) fitting the model to (sampled) observations to minimise variational free energy (*F*; surprisal)(Friston *et al.*, 2017a; Friston, 2019; Sajid *et al.*, 2019) and, 2) selecting actions that minimise expected free energy (*G*; uncertainty)(Parr and Friston, 2018; Da Costa *et al.*, 2020). The variational free energy is the complexity cost incurred in forming accurate posterior beliefs about causes of sensation:

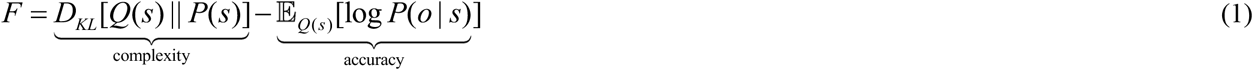

and, expected free energy:

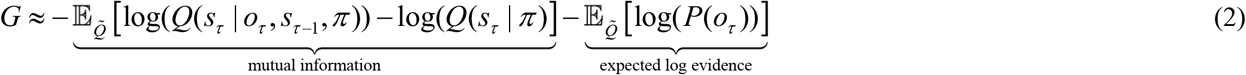

Where 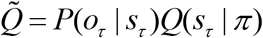 and *Q*(*o*_*τ*_ | *s*_*τ*_, *π*) = *P*(*o*_*τ*_, | *s*_*τ*_).

Using these objectives, expectations about hidden states, policies and precision are optimized through inference, and model parameters optimized through learning. This involves the variational message passing of sufficient statistics of posterior beliefs (i.e., expected probability) among neuronal populations. Note, the variational message passing can be formulated as a gradient descent on variational free energy, using a mean-field approximation (Friston *et al.*, 2017d; Parr *et al.*, 2019a).

### 2.1 Deep temporal models

The process theory underwriting active inference is based on a partially observable Markov decision process (POMDP). This can be defined as a generative model with discrete outcomes that are caused by discrete hidden states (Friston, FitzGerald et al. 2017, Friston, Lin et al. 2017). These models can have a deep temporal (i.e., hierarchical) structure, where the outcomes of one level generate the hidden states at the level below (Friston *et al.*, 2017c; Friston *et al.*, 2017d). The equation below defines the factorized form of a deep temporal model, via its joint probability over outcomes and hidden states.

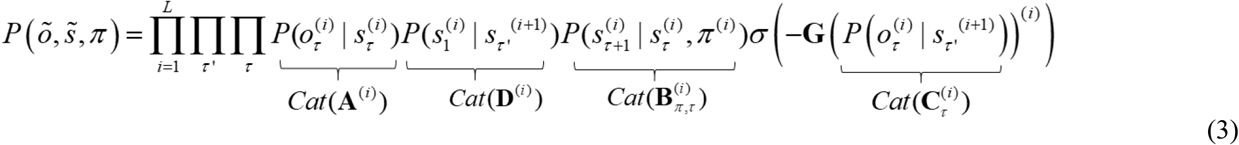

where *õ* and 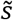 denote sequences of outcomes and hidden states respectively, until the current time point: *i* denotes the *i*-th hierarchical level and *L* the total number of levels and policies, *π* are trajectories over potential action space.

Here, outcomes depend upon hidden states and hidden states depend upon policies. Outcomes and hidden states are parametrized by two distinct categorical distributions: *A*^(*i*)^ and 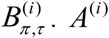 is the likelihood function, that maps hidden states to outcomes at the *i*-th level, and 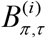 the distribution that maps transitions from one hidden state to the next, at the *i*-th level, under policy *π*. Successive levels of the generative model are linked by *D*^(*i*)^, which defines the mapping between hidden states at *i+1-*th level to the initial states at the *i*-th level below.

Policies, and the next action, are selected by sampling from the softmax function of expected free energy. Consequently, policies are more probable, *a priori*, if they minimise expected free energy. Using this, hidden state sequences are generated using 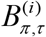 determined by the selected policy. These hidden states generate outcomes and initial hidden states in the level below (according to *A*^(*i*)^ and *D*^(*i*)^). They influence the expected free energy through *C*^(*i*)^ and the policies that determine transitions among subordinate states. Here, the model parameters, i.e., 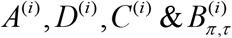 are equipped with a prior (categorical) distribution. The key aspect of this generative model is that transitions among states proceed at different rates at different levels of the hierarchy. Essentially, hidden states at higher levels contextualize trajectories of hidden states at lower levels, enabling a deep dynamic narrative (Friston *et al.*, 2017d).

This generative model can be regarded as the ‘structure’ in structure-function relationships and is thought to underwrite functional brain architectures that realize active inference. This perspective allows us to associate *in-silico* lesions with anatomical lesions. Specifically, when defining model inversion in terms of message passing, *A*^(*i*)^ can be regarded as extrinsic connections i.e., between different regions/neuronal populations while *B*^(*i*)^ can take the form of intrinsic connections i.e., within and between the cortical layers of a single region(Parr *et al.*, 2019b).

### 2.2 Learning

Each synthetic subject has implicit prior beliefs about their model parameters (i.e., likelihood - *A*^(*i*)^, transitions - *B*^(*i*)^, etc). This includes prior beliefs over model parameter priors (i.e., hyperpriors), which are learned through Bayesian belief-updating (Friston *et al.*, 2017a; Friston *et al.*, 2017b). The natural choice for the conjugate hyperprior over categorical priors is a Dirichlet distribution. This means that hyperpriors can be expressed simply in terms of Dirichlet concentration parameters (Parr, 2019), which represent each state-to-outcome (for *A*^(*i*)^) and state-to-state (*B*^(*i*)^) mappings or connectivity:

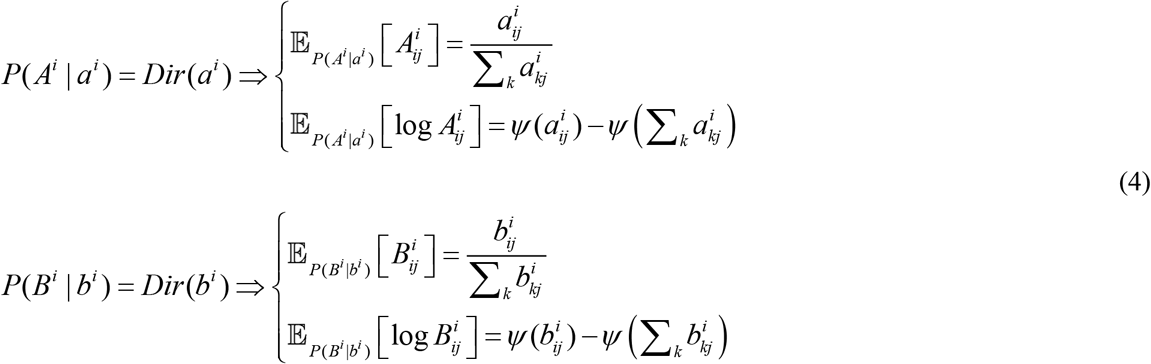

where *ψ* is the digamma function.

The Dirichlet parameters can be thought of as ‘pseudo-counts’ i.e., as observations are made, they accumulate Dirichlet parameters that best assimilate sensory data. The more often a given pairing (of state and outcome, or past and present state) is observed, the greater the number of counts attributed to that pairing (Beal, 2003; Blei *et al.*, 2003). Thus, beliefs change to a greater extent when the counts are low—because few pairings have been observed—than when they are high. This accumulation process closely resembles Hebbian plasticity (Hebb, 1949), where synaptic efficacy is reinforced upon the simultaneous firing of a pre and postsynaptic neuron (Friston *et al.*, 2016). This learning process affords experience-dependent plasticity: namely, the strengthening of synaptic connections during belief updating.

### 2.3 Simulating *in-silico* lesions

*In-silico* lesions can be simulated by manipulating precision, where precision scores confidence (i.e., the inverse of uncertainty) in beliefs about the causes of sensations. As in our prior work (Sajid *et al.*, 2020b; Sajid *et al.*, 2020c), we manipulate precision, over model parameters *A*^(*i*)^ and *B*^(*i*)^ which results in different types of damage that can be linked to pathological lesions in the human brain.

Precision over *A*^(*i*)^ (sensory precision) corresponds to the confidence with which the model can infer the cause of observations on the basis of prior experience. Decreasing the precision over *A*^(*i*)^ mimics a lesion to extrinsic connections (i.e., between regions expectations about causes and observations). This results in uncertainty about the causes of observations. Extending our prior work(Sajid *et al.*, 2020c), we distinguish between structural and functional disconnections. When precision over *A*^(*i*)^ is completely imprecise (i.e., probability distribution is uniform) we induce a “structural disconnection” between the observations and their underlying causes, meaning that it is not possible to resolve the causes. A less severe lesion (i.e., slightly imprecise distribution), in contrast, only results in a “functional disconnection” leaving imprecise, ambiguous relationships between causes and outcomes, that can, in principle, be resolved through re-learning.

Precision over *B*^(*i*)^ (state transition precision) corresponds to confidence with which the model can predict the present from the past (i.e., infer state transitions using prior transition probabilities). Decreasing precision over *B*^(*i*)^ induces a focal lesion to intrinsic connections (e.g., within and between the cortical layers of a single region). This precision affects the model’s ability to learn (i.e., limiting the ability to update beliefs about causes). As with lesions to precision over *A*^(*i*)^, lesions to precision over *B*^(*i*)^ can either be structural (completely imprecise) or functional (slightly imprecise). Structural lesions prevent appropriate learning and belief updating. Functional lesions reduce the capacity to learn and update beliefs.

Belief updating is also compromised when sensory precision (i.e., over *A*^(*i*)^) is reduced. This can be linked to neuromodulatory control when precision is implicated at the lower level in deep temporal models. In other words, the higher level must appropriately disambiguate between similar competing lower-level causes and modulate their precision. For example, acetylcholine release, in extrinsic connectivity, is thought to increase sensory precision (e.g., by boosting bottom-up sensory signals) enabling the brain to respond optimally(Moran *et al.*, 2013). Lowering precision over *A*^(1)^ is therefore expected to impair neuromodulatory control of synaptic efficacy in intrinsic connections.

### 2.4 Simulating electrophysiological measurements

The form of the (variational) message passing mandated by active inference, as shown below, allows us to associate variables with idealised electrophysiological measurements (Friston *et al.*, 2017a; Parr *et al.*, 2019b):

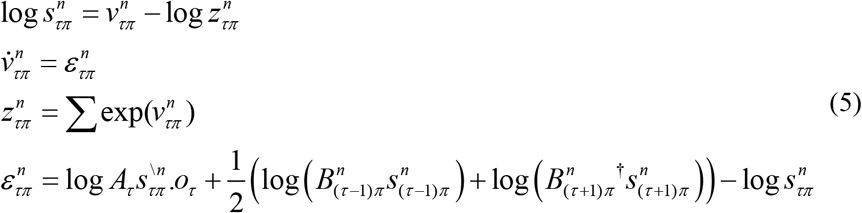

Here, 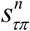 denotes the expected hidden state factor (*n*) at time (*τ*), conditioned on a policy (*π*); *z* is the partition function. *v* and *ε* are auxiliary variables that play the roles of membrane depolarization (i.e., post-synaptic potential) and prediction error, respectively. *v* is computed from the inputs of other neurons and transformed to *s* via the partition function, *z*. Note, that *s* is the signal that is propagated to other neuronal populations and is analogous to firing rate, as measured by single unit recordings. The rate of change of *v* can be associated with local field potentials, after bandpass filtering between 4 Hz and 32 Hz (Friston *et al.*, 2017a).

## 3. A Generative Model of Word Repetition

To illustrate how particular lesions could trigger different recovery mechanisms, we extended our previous generative model (Sajid *et al.*, 2020b; Sajid *et al.*, 2020c) and active inference scheme for simulating word repetition (Ueno *et al.*, 2011; Moritz-Gasser and Duffau, 2013; Nozari and Dell, 2013). The subject (i.e., model) hears a single spoken word and must repeat it. If repeated correctly, they receive positive feedback (and negative otherwise).

### 3.1 Previous model of word repetition

In the previous version of the model, there were three state factors (Target Word, Spoken Word, Epoch) and three outcome modalities (Proprioception, Word, and Evaluation). The Target Word factor has five states, corresponding to the words that could be heard: red, blue, table, triangle and square. The Spoken Word factor contains beliefs about what word should be repeated: red, blue, table, triangle and square. The Epoch factor codes two different stages of the trial: listening to a target word (epoch 1), repeating it and receiving evaluation (epoch 2). In terms of outcomes, the Proprioception outcome reports whether the subject moved their mouths to speak or not. This depends on the state of the epoch factor: if the subject believes they are in epoch 1 (listening to the target word), then they are not speaking but if they believe they are in epoch 2 (repeating the target word), then they are speaking. The Word outcome reports the heard word (target or spoken): red, blue, table, triangle or square. This depends on the states of the Target Word, Spoken Word and Epoch factors. If the subject believes they are in epoch 1, the word outcomes depend on the Target Word factor but if the subject believes they are in epoch 2, the word outcomes depend on the Spoken Word factor. Finally, the Evaluation outcome indicates if the spoken word was the same as the target word. The outcome depends on the Epoch factor: positive (correct) if the spoken word is the same as the target word at epoch 2, negative (incorrect) if the spoken word is not the same as the target word at epoch 2, or neutral (during epoch 1). For a discussion of the functional architecture entailed by this generative model—and the underlying neurobiology—please see Sajid, et al (2020)(Sajid *et al.*, 2020c).

### 3.2 Current (hierarchical) model of word repetition

The current model extended our previous model (described above) in two ways. First, the new model is hierarchical with two levels (Figure 1). The top level (not included previously) has one state factor: Network. The Network factor contains two states: premorbid or alternative. The premorbid system is associated with greater precision than the alternative system—as if the model were in an attentive compared to an inattentive state, respectively. This follows because changes in precision are generally associated with attention (Brown *et al.*, 2011; Parr and Friston, 2017).

**Figure 1.**
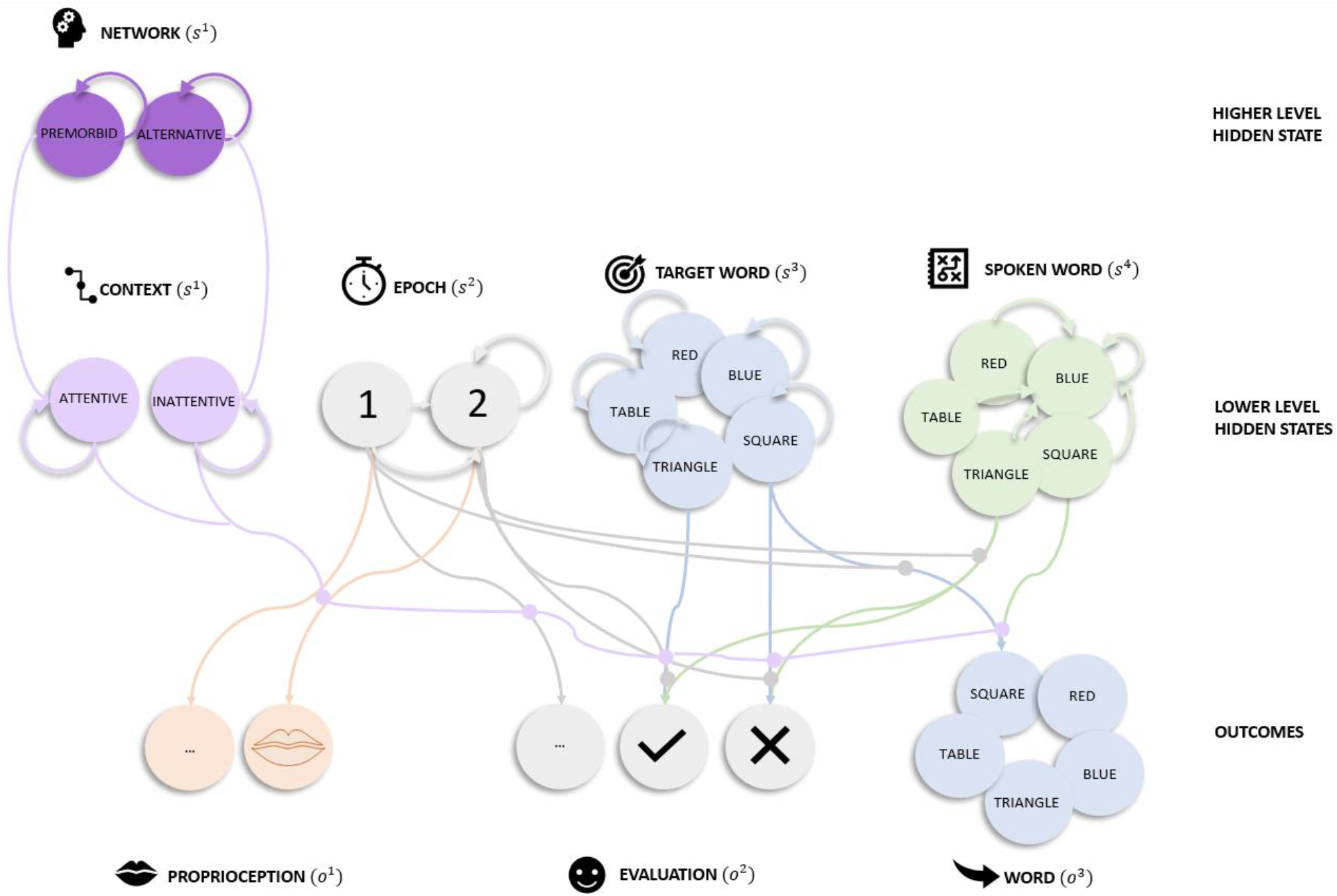
(Generative Model) Graphical representation of the generative model for word repetition. The model comprises two levels. There is one higher-level state factor (Network), four lower-level state factors (Epoch, Target Word, Spoken Word, and Context) and three outcome modalities (Proprioception, Evaluation and Word). *Network* (2 states) at the higher level specifies the system in play (premorbid or alternative). *Epoch* (2 states) indexes the phase of the trial. During the first epoch, the target word is heard. The second epoch involves repeating the word and concomitant evaluation. A positive evaluation is provided if the Word outcome matches the Target Word state, and a negative evaluation otherwise. The *Target Word* factor (5 states) lists the words the experimenter can ask the participant to repeat. The *Spoken Word* factor includes the words that the model can choose to say (5 states). The *Context* factor (2 states) specifies whether the subject is in an attentive or inattentive state. Lines from states to outcomes represent the likelihood mappings and lines mapping states within a factor represent allowable state transitions. The lines represent plausible connections, and their absence reflects implausible connections. To avoid visual clutter, only a few likelihoods and transition probabilities are shown, but they are consistent across the different state factors and outcome modalities. For example, the Word likelihood mapping from the Target Word state (square) to the Word outcome (square) is shown for Epoch 1, but similar mappings apply when mapping between the blue Target Word and blue Word outcome and between the triangle Target Word state and triangle Word outcome, etc. One (from a total of 5) example transition probability is highlighted for the Spoken Word state, i.e., the transition is always to blue, regardless of previously spoken word (red, table, triangle, square or blue). This transition represents the choice to say ‘blue’. Similar mappings are applied when choosing to say ‘triangle’. Actions then correspond to the selection of particular transition probabilities. The Context factor modulates the strengths of the likelihood mappings between the other factors and outcomes, depending on whether the model is in an attentive or inattentive state.

The second extension was the addition of a fourth state factor (Context) to the lower level that serves as the target of top-down messages from the hierarchical level above. The Context factor denotes attentive (precise) or inattentive (imprecise) beliefs depending on the Network factor at the higher level: If the subject believes they are using the premorbid system, then they will be in the attentive context, and otherwise inattentive. A shift in context from attentive to inattentive at the lower level reduces the precision of the mappings between other state factors and outcomes i.e., decreased confidence about the causes of sensations. Notice that this deep model equips our synthetic subject with a more nuanced and context sensitive processing capacity that can be likened to having an attentional set, which is sensitive to—and contextualises—processing at lower levels.

The context factor is particularly important for the current simulations because it effectively duplicates the message passing under the remaining factors (c.f., premorbid and alternative systems). These context states are distinguished only in terms of the sensory precision in the mappings between states and outcomes. Therefore, when precision in these mappings is similar for both contexts, the higher (Network) level cannot infer the appropriate attentional set or context. The Network factor at the higher level is only connected to the Context factor at the lower level. This setup enables beliefs about the lower-level Context factor to be updated if there is a change in beliefs about the Network at the higher level. All of the other state factors at the lower level are conditionally independent of the higher level—and are exactly the same as on our previous model.

### 3.3 Model parameters

Figure 1 illustrates how States are connected to Outcomes using lines or edges. The strengths of these connections are defined by the likelihood parameters, *A*^(*i*)^ which indicate the sensory precision of extrinsic connections (Section 2.3 above). Each state also has a transition matrix, *B*^(*i*)^ (denoted by arrows in Figure 1) that maps the state at the current timepoint to the next time point. This indicates how precisely the current state will transition to another state (via intrinsic connectivity).

For the Spoken Word factor, there are five transitions. These transitions depend upon the action that the subject selects: upon taking an action, this state always transitions to the selected word. For example, when the subject chooses to say ‘blue’, regardless of previous the word (i.e., red, triangle, etc), the Spoken Word state will be blue (highlighted in Figure 1). For the lower-level Epoch factor, states transition from epoch 1 to epoch 2 which is an absorbing state (final epoch). For the Context factor and the Target Word factor, transitions were represented in an identity matrix (of size two and five). This means these factors stay the same throughout a trial. The transitions were also represented with an identity matrix for the higher-level network factor.

The model had strong preferences for receiving positive evaluation (i.e., getting the repetition correct) and strong aversion to negative evaluations. It was also equipped with 5 different one-step-ahead polices (action trajectory) to choose from, corresponding to each word that could be spoken. We specified the following hyper-parameters: threshold for passing back control to the higher-level was *e*^−16^ ≈ 0 and the decaying learning rate was 2.

### 3.4 Relationship to other models

The word repetition model was implemented as a partially observable Markov decision process and some parallels can be drawn between this and existing models. Our model has two parallel contexts, defined by the higher-level state factor, which specifies a premorbid and alternative system. This equips the model with two-way (albeit asymmetric) processing, due to imbalanced (attentional) resource allocation, that may be involved in word repetition. The asymmetry here is that the subject gives prior preference to attentive (i.e., precise), over inattentive (i.e., imprecise) beliefs. In contrast, a symmetrical model would select attentive or inattentive states, with equal probability. This makes our model formally similar to the recurrent neural network presented in (Chang and Lambon-Ralph, 2020), with dual structures that support function, but are asymmetrical due to differences in computational resource allocation.

Our Bayesian approach is formally distinct from some previous (cognitive) models of word repetition; for example, recurrent (Ueno *et al.*, 2011; Chang and Lambon-Ralph, 2020) and adaptive (Tourville and Guenther, 2011; Guenther and Vladusich, 2012) neural networks. These formulations can be regarded as deterministic function approximators, after an initial training phase. In contrast, our approach does not require ‘training’ and can be used to simulate outcomes based on a (learnt or prespecified) probability distribution or belief state.

Our model has close ties to the state feedback control model of speech production using Kalman filtering, another Bayesian algorithm (Houde and Nagarajan, 2011). The Kalman filter is the Bayes-optimal solution under a linear generative model, but a cascade of such solutions would not be the optimal solution to (non-linear and non-Markovian) hierarchical models (Perrinet *et al.*, 2014). Conversely, active inference considers the hierarchical system as a whole and provides an optimal solution in the form of variational (Bayesian) filtering or belief updating.

## 4. Simulating the effect of in-silico lesions to our word repetition model

### 4.1 Control simulation (Lesion 0)

To measure differences in recovery, we simulated a control model without any lesion. This acts as a sanity check for the architecture of the premorbid system and simulates the behaviour of healthy subjects performing the task. At the top level, the premorbid system was given greater precision than the alternative system (see Lesion 0, Table 1). This increased attention to the premorbid system at the lower level(Brown *et al.*, 2011; Parr and Friston, 2017) resulting in strong prior expectations to use the accompanying premorbid system. To prevent re-learning of model parameters, the Dirichlet concentration parameters (for intrinsic and extrinsic connections) were set at high values— as if the subject had over-learned the task (see Lesion 0, Table 1).

**Table 1.** (Overview of precision manipulation) The table presents a breakdown of the five models and the underlying changes in precision that simulated particular lesion types—and the initial Dirichlet counts. 1 denotes high precision, 0 low precision (i.e., a uniform distribution) and everything else a gradation between the two.

| Lesion | Sensory precision $A^1$<br>in<br>premorbid system | Sensory precision $A^1$<br>in<br>alternative system | Target word<br>transition $B^1$<br>precision | Dirichlet<br>Count |
| --- | --- | --- | --- | --- |
| 0 | 1.0 | 0.9 | 1.0 | 1000 |
| 1 | 0.0 | 0.9 | 1.0 | 1000 |
| 2 | 0.0 | 0.9 | 0.4 | 1 |
| 3 | 0.1 | 0.1 | 1.0 | 1000 |
| 4 | 0.1 | 0.1 | 0.4 | 1 |

### 4.2 Disabling the premorbid system and engaging the alternative system (Lesion 1)

Within the premorbid (attentive) system, we introduced structural disconnections to the extrinsic connections linking state factors (Target Words, Spoken Words and Epoch) to outcomes (Proprioception, Words and Evaluation), see Figure 1 and Table 1, Lesion 1). Now the subject is unable to use the premorbid system to differentiate between implausible and plausible hidden states because there is no evidence for any particular Target Word causing the Word outcome—and no systematic mapping between the Spoken Word and Evaluation. In light of this, the disconnected premorbid system is also unable to learn anything. In contrast, all the extrinsic connections in the alternative (inattentive) system were preserved. As in the control model (Lesion 0), re-learning of the model parameters in the alternative system was precluded with high Dirichlet concentration parameters (for both intrinsic and extrinsic connections).

Despite structural disconnections to the premorbid system, and a strong prior expectation at the higher level to use the premorbid state (as in the control model), we expect that Lesion 1 will be able to perform word repetition without any re-learning. This is because the alternative system can support the same function—albeit with lower precision. Functional re-organisation is still required, however, because the lesioned subject still needs to (i) correctly infer (at the higher level) that there has been a change in the options available (since the premorbid option has been rendered ineffective for the task at hand) and (ii) shift, at the higher level, from using the (damaged) premorbid system to the intact alternative system. This would be evidenced by a change in beliefs at the higher level of the model. If the subject fails to engage the alternative system, incorrect responses will be produced. In brief, any recovery depends on rapid updates to higher-level beliefs about the options available for task performance. This should happen after a few trials, when the model gets incorrect feedback, when attempting to use the premorbid system.

### 4.3 Long-term experience-dependent plasticity in the alternative system (Lesion 2)

The second intervention was the same as the first, but with an additional functional lesion to the intrinsic connections mediating state transitions, *B*^1^, for the Target Word factor (see Figure 1 and Lesion 2 in Table 1). We expected this to increase prediction errors when inferring the target word at Epoch 2.

In addition to (i) the structural lesions to extrinsic connections in the premorbid system and (ii) functional lesions to intrinsic connections which impacts on the alternative system, we also (iii) reduced the Dirichlet parameters (for intrinsic and extrinsic connections). Now the subject can accumulate Dirichlet parameters that best account for experiences (via future learning) thereby strengthening synaptic connections of model parameter *B*^1^ (Friston *et al.*, 2016; Da Costa *et al.*, 2020) in the alternative system. This is consistent with experience-dependent plasticity. In brief, we expected that the subject would partially recover slowly through experience-dependent plasticity within the alternative system.

Other forms of *in-silico* intrinsic lesions would have impeded the model’s ability to exhibit adaptive experience-dependent plasticity. For example, introduction of mis-specified or jumbled (as opposed to imprecise) Dirichlet priors would have resulted in maladaptive learning of Dirichlet priors over the course of the simulations.

### 4.4 Manipulating the neuromodulatory balance between the premorbid and alternative systems (Lesion 3)

To disrupt the neuromodulatory balance between the premorbid and alternative system, we induced functional lesions to the likelihood mappings for extrinsic connections to the Word, Proprioception and Evaluation outcomes, see Figure 1 and up 3 in Table 1. Compared to Lesion 1, the premorbid system is not completely disconnected, and the alternative system is less precise. Now the subject can use both systems (albeit ineffectively) but no longer has the appropriate machinery to control which system to use. In addition, as in Lesion 0 and 1, we prevented re-learning of model parameters by using high Dirichlet concentration parameters (for intrinsic and extrinsic connections) in both systems (see Lesion 3 in Table 1).

### 4.5 Multifocal lesions (Lesion 4)

In Lesion 4, we repeated Lesion 3 (functional lesions to extrinsic mappings in premorbid and alternative systems) and additionally lesioned the intrinsic connections as in Lesion 2, (see Table 1). These multifocal lesions were expected to further impede functional recovery relative to all prior models. Nevertheless, the system retained learning capacity because the Dirichlet concentration parameters were set to be low (see Lesion 4, Table 1).

## 5. Results of simulations

For each lesion, we evaluated word repetition performance (Figure 2) and lesion severity (Figure 3 & 4) over 1000 trials. Lesion severity is quantified using: (i) free energy, (ii) degeneracy and (iii) redundancy. As a reminder, free energy is the complexity cost incurred in forming accurate posterior beliefs about the causes of sensations/outcomes. As policies are more probable, *a priori*, if they minimise the expected free energy, model evidence is higher when free energy is lower. Degeneracy, in this formal setting, is the many-to-one mapping between hidden causes and outcomes (i.e., entropy of posterior beliefs). This has the advantage of providing flexibility in our internal explanations for sensory outcomes (keeping our options open, allowing re-learning) but can also result in less confidence (more uncertainty) in what is causing the outcomes (lowering accuracy). Finally, redundancy is the inverse of efficiency and represents the complexity cost of updating beliefs when there is multiple alternative (i.e., degenerate) causes to consider (i.e., more belief updating is required).

**Figure 2.**
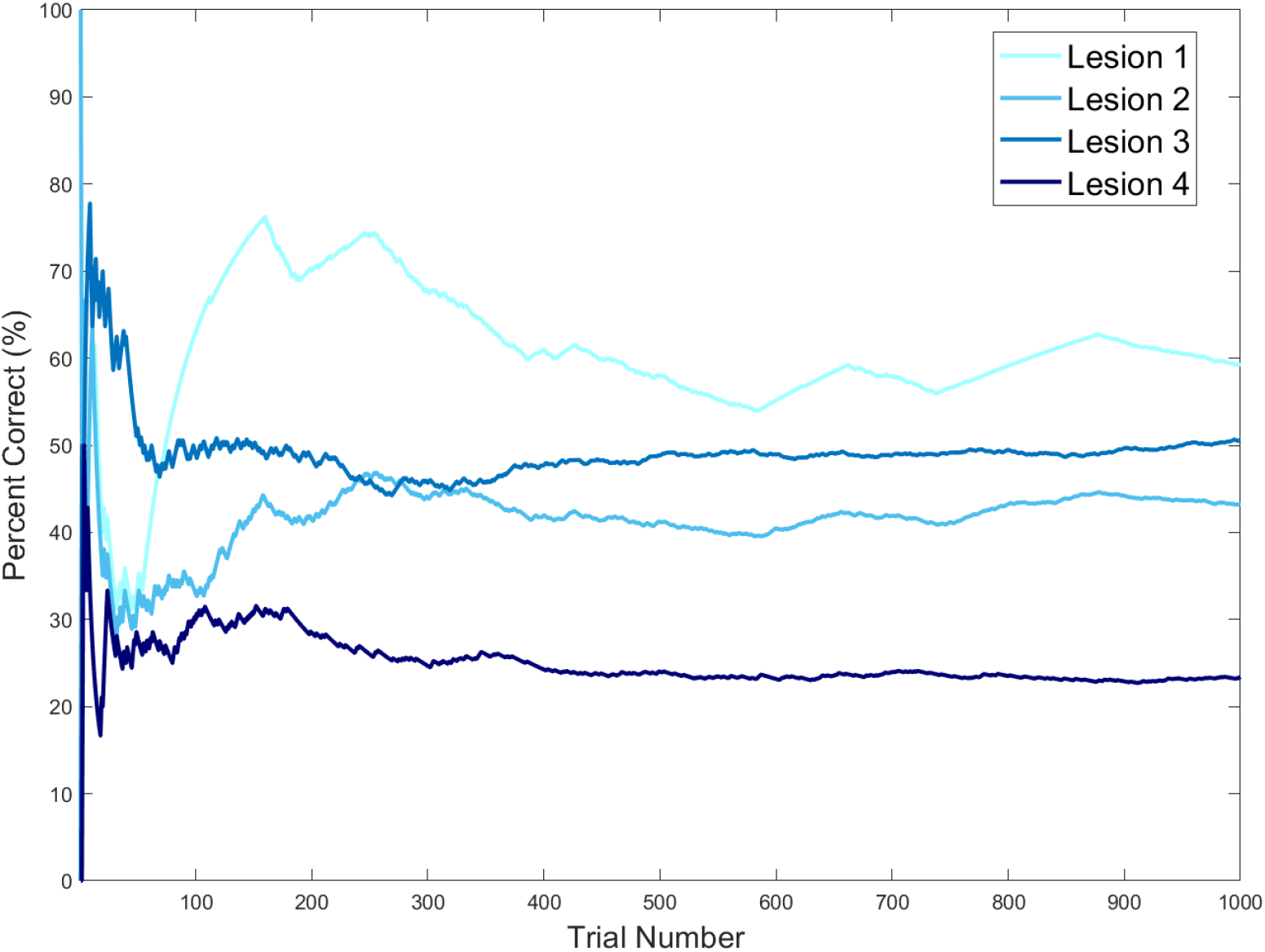
(Behavioural performance across 1000 trials) This plot shows the percentage of correct responses for each model across 1000 trials. The x-axis is the trial number, and the y-axis is the cumulative percentage of correct responses (i.e., the percentage of trials that were correct for all trials up to the current trial number).

**Figure 3.**
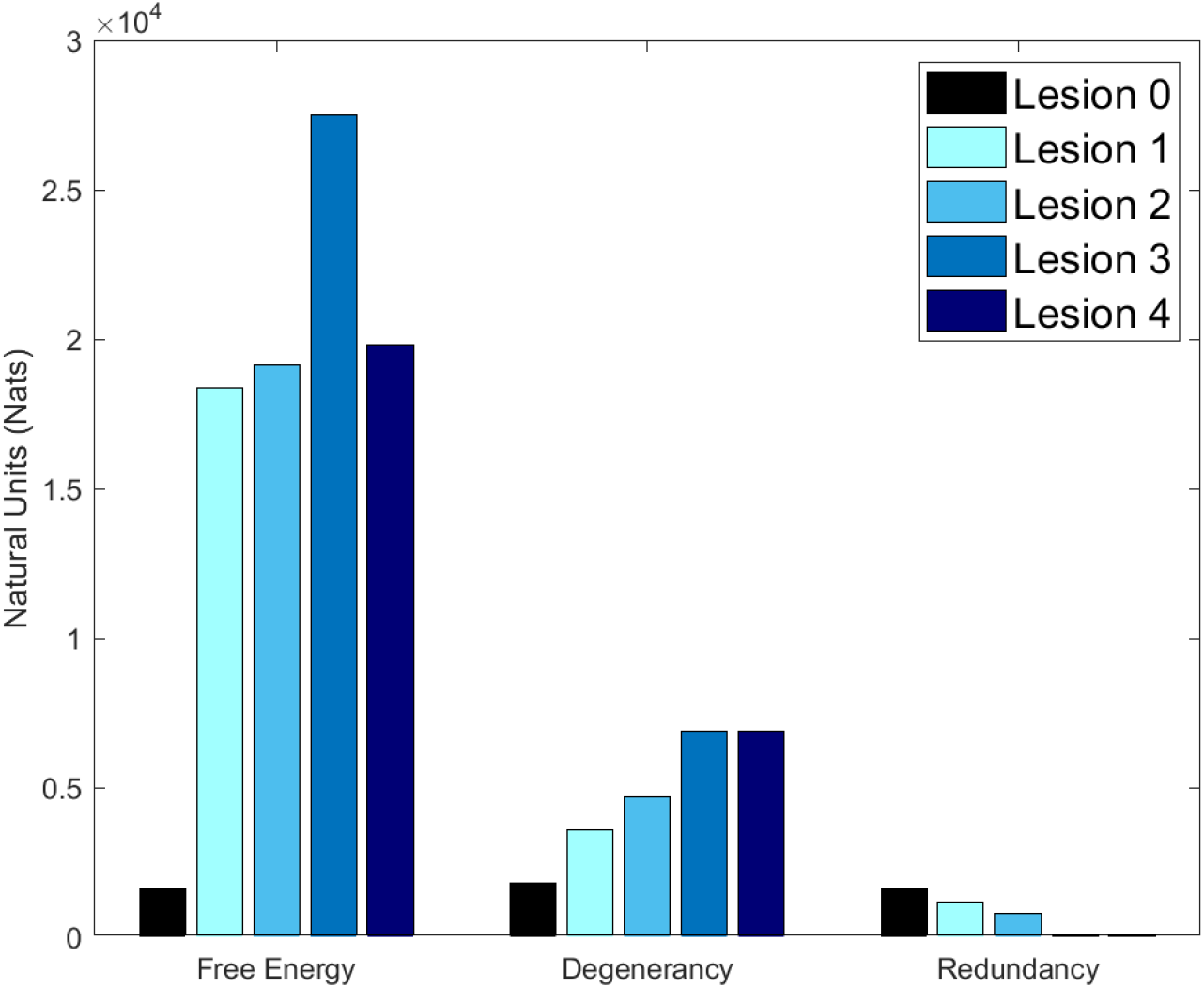
(Free energy, degeneracy, and redundancy) This bar plot reports the total free energy, degeneracy, and redundancy for each kind of lesion across 1000 trials. Free energy is the model evidence, degeneracy is the entropy of posterior beliefs about the causes of sensations and redundancy is the complexity cost incurred by forming those beliefs. The y-axis represents information, measured in natural units. Both Lesion 3 and Lesion 4 have redundancy < 30 nats – due to an inability to update posterior beliefs away from prior beliefs.

**Figure 4.**
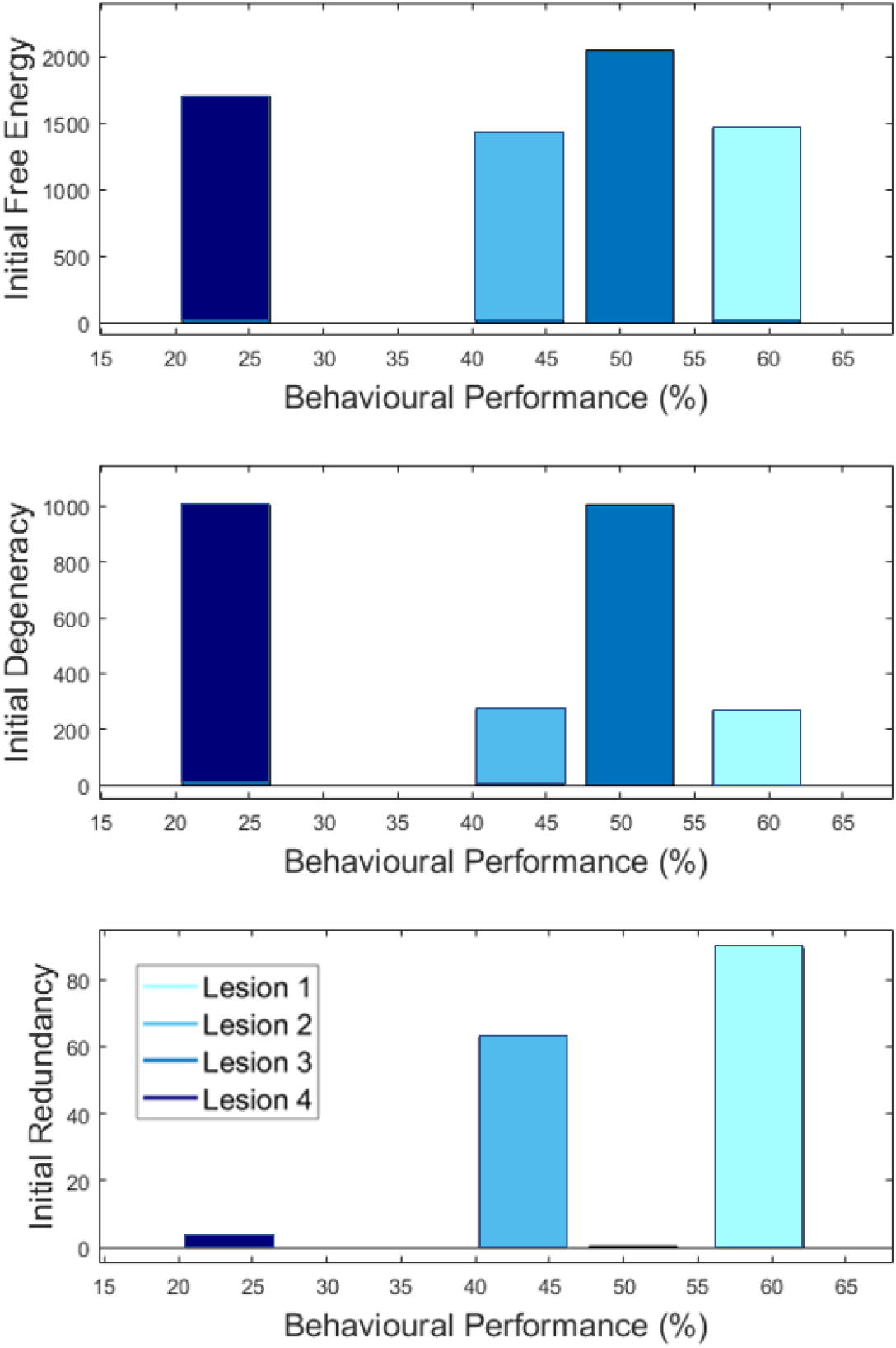
(Lesion severity and behavioural performance). For each model, the plots report (in nats) free energy (top row), degeneracy (middle row) and redundancy (bottom row), accumulated over the first 50 trials (y axis), according to behavioural performance (percent correct responses measured after 1000 trials in x axis).

### 5.1 Control simulation (Lesion 0)

In the absence of lesions, our synthetic subject was able to repeat the correct word with 100% accuracy across all 1000 trials. This indicates that the subject correctly inferred that they were operating in an ‘attentive’ (i.e., precise) context, relying on the premorbid system, despite being equipped with an alternative system.

### 5.2 Disabling the premorbid system and engaging the alternative system (Lesion 1)

After a sharp drop in performance in the first 50 trials, Lesion 1 rapidly regained performance, with an early performance peak (76% around Trial 160), and stabilisation to ~58% by the 400^th^ trial that persisted across the remaining trials (Figure 2; cyan line). Overall performance for this model was 60%.

Compared to Lesion 0 (the control model), Lesion 1 had lower (i) accuracy; (ii) model evidence (high free energy) and (iii) degeneracy (representational flexibility) because, when the premorbid system was unavailable, the alternative system (with less precise likelihood mappings) was engaged. The novel insight here is that Lesion 1 was able to shift context (to attend to the alternative system) even though the network level indicates that the premorbid system should be used. This models a higher-level reorganisation of the neural circuitry involved in repeating words that arose naturally because the premorbid system was damaged and was therefore unable to supply precise evidence to the higher level.

### 5.3 Long-term experience-dependent plasticity in the alternative system (Lesion 2)

Lesion 2 had impaired behavioural performance, across trials, relative to Lesion 1 (43% versus 60%) reflecting the additional lesion to intrinsic connections in the alternative system that compromised new learning. The residual learning capacity in Lesion 2 is illustrated by observing the change in Dirichlet parameters over time (Figure 5). This shows how Lesion 2 slowly shifts towards the ‘optimal’ (Lesion 0) distribution, with saturation effects.

**Figure 5.**
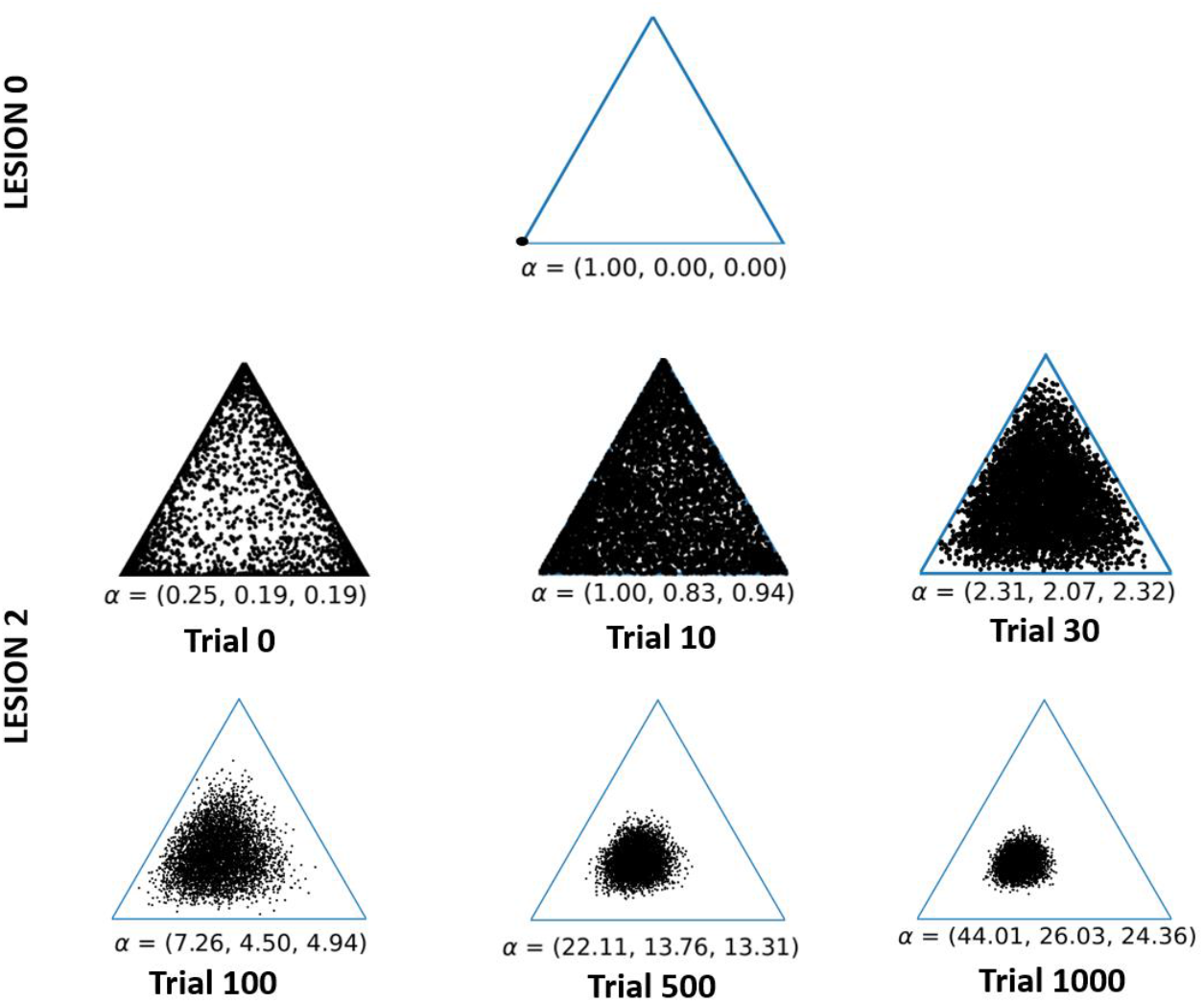
(Synaptic strength changes via experience-dependent plasticity) The figure illustrates the Dirichlet distribution in a 3-dimensional coordinate space, i.e., 2-simplex. The concentration of dots in one corner reflect precise beliefs; and scattered dots denote imprecise beliefs. Each dot represents a single sample from the Dirichlet distribution (determined by the alpha parameters), and each plot displays 5000 samples. For visual clarity, we have focused on the changes in the Target Word transitions for how the word ‘red’ can potentially transition to either ‘red’, ‘blue’ or ‘table’. Similar distributional shifts (i.e., learning) apply to other Target Word transitions. The plot for the control model illustrates that, with a precise transition mapping, the distribution is highly concentrated at one corner of the plot (to the extent that the points look like a single dot). On the second row, a series of panels illustrate the learning process for Lesion 2 over six time points (trials 0, 10, 30,100, 500 and 1000). Immediately following the lesion (trial 0) a scattered distribution is evident, despite a higher concentration of points at all corners of the triangle. However, as the lesioned model learns (between trials 0 and 1000), the distribution converges to the corner associated with the high alpha—more closely resembling the control.

### 5.4 Manipulating the neuromodulatory balance between the premorbid and alternative systems (Lesion 3)

In the first 50 trials, Lesion 3 performed better than Lesion 1 because the premorbid system retained some capacity (was not completely disconnected). However, unlike Lesion 1, the performance of Lesion 3 did not improve over time, stabilising at about 50%, after trial 200 (see Figure 2). This is because (i) Lesion 3 struggled to infer which system (premorbid or alternative) was appropriate and (ii) both the premorbid and alternative systems were generating errors because precision in the extrinsic connections was low. Consequently, compared to Lesions 1 and 2, Lesion 3 had higher free energy (less model evidence) and higher degeneracy (more uncertainty) across trials. This lesion also produced lower redundancy that reflects an inability to appropriately update posterior beliefs i.e., the model can no longer estimate which causes were responsible for the outcomes.

### 5.5 Multifocal lesions (Lesion 4)

Lesion 4 performed worse than Lesion 3, across trials, demonstrating the impact of compromised learning. It’s also interesting to note that Lesion 4 had less free energy than Lesion 3 (i.e. greater model evidence) because it had higher levels of redundancy and accuracy (free energy = redundancy – accuracy)(Sajid *et al.*, 2020c).

## 6. Physiological predictions

Recovery has previously been associated with various physiological changes e.g., increased baseline firing frequency (Schiene *et al.*, 1996; Carmichael, 2003), synchronous neural activity (Carmichael, 2003), etc. The *in-silico* lesions reflect these changes in neuronal functionality due to missing/interrupted circuitry (via degeneracy) and alterations within surviving structures (via peri-lesional activity) that affect the belief updating process.

The lesioned models demonstrate reduced baseline evoked response frequency and a shifted inhibitory evoked response, relative to control (Figure 6). Lesion 1 has similar evoked response magnitude to control, but a decreased presynaptic excitatory potential across all trials (Figure 6; second row) – similar trends have been observed in aphasic subjects during picture naming tasks (Laganaro *et al.*, 2008). Additionally, the initial trials exhibit an attenuated evoked response, but later trials expose a steady increase in evoked responses (Laganaro *et al.*, 2008). Lesion 2 has an attenuated, but noticeable, presynaptic inhibitory potential (Figure 6; third row) and muted overall excitatory potential. Lesion 3 has the highest decrease in evoked response, compared to other models, alongside an decrease in postsynaptic inhibitory potential (Figure 6; fourth row)(Kotz and Friederici, 2003). Muted evoked response can be observed for Lesion 4 (Figure 6; fifth row).

**Figure 6.**
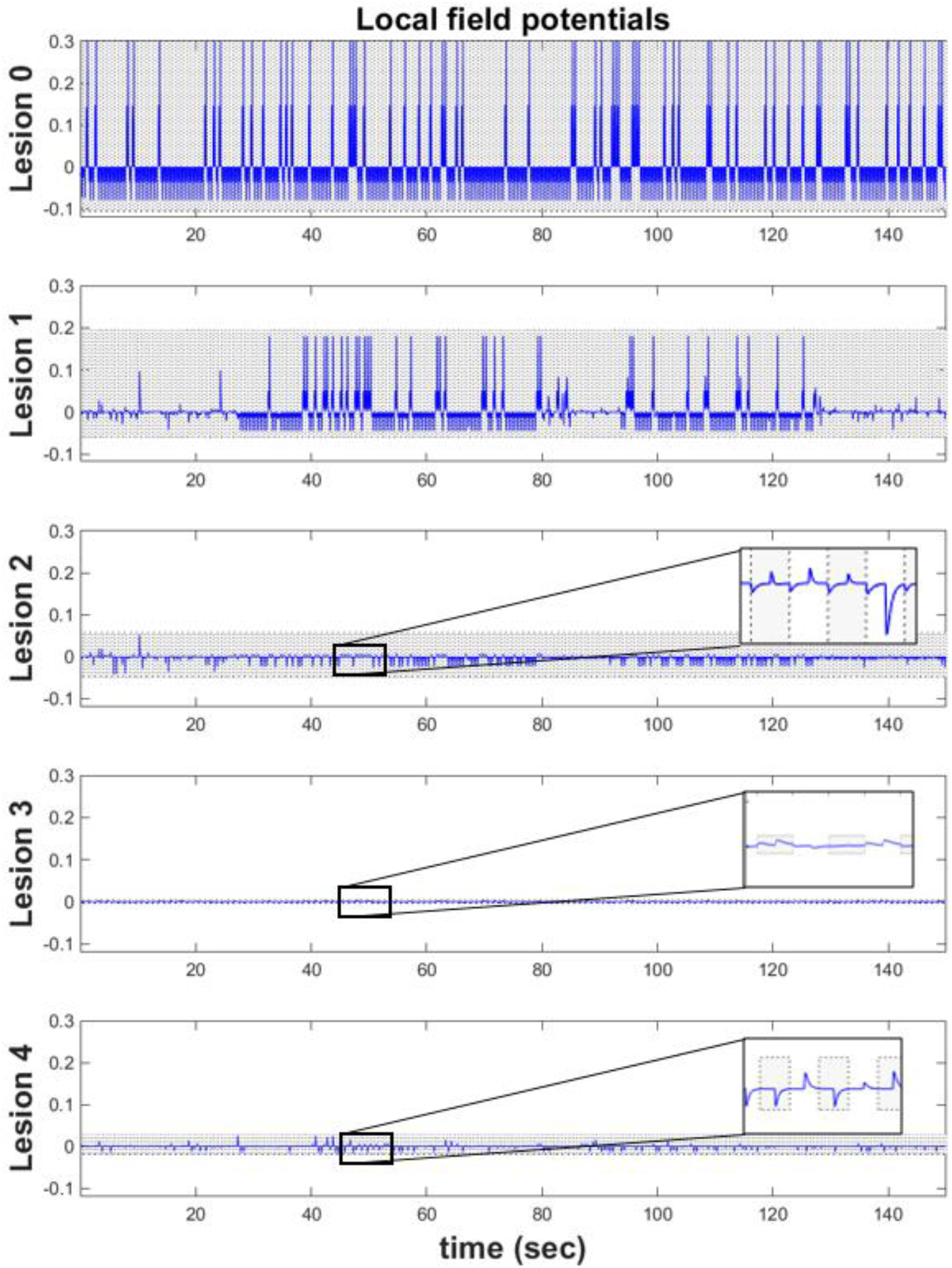
(Simulated local field potentials) These plots show the temporal changed in simulated local field potentials for the Target Word ‘table’ neuronal population across the first 300 trials. The blue line represents the trajectory of evoked responses for the Target Word ‘table’ over some arbitrary unites (y-axis). The simulated lesion models are presented in the following order: Lesion 0, Lesion 1, Lesion 2, Lesion 3 and Lesion 4.

This simulated electrophysiology serves three key purposes. Firstly, it lends a construct validity to the generative model for word repetition (and the inferential process); these simulations are congruent with real electrophysiological measurements from humans (Kotz and Friederici, 2003; Laganaro *et al.*, 2008; Pei *et al.*, 2011) and other primates (Luhmann *et al.*, 1995; Schiene *et al.*, 1996). Second, it offers an intuition about the computational processes that might underpin these responses, and how they change when particular processes are perturbed. Finally, we can derive quantitative predictions about measured electrophysiology from human subjects and use them to quantify recovery mechanisms. Our simulations suggest that different perturbations lead to different neuronal responses, so it should be possible—in principle—to use electrophysiological data to discriminate between different types of impairment. However, further work is required to anatomically ground our word repetition models.

## 7. Discussion

In this study, we developed a generative model that is equipped with two distinct systems for independently performing a word repetition task. One system has high sensory precision allowing it to reproduce attentive and accurate behaviour. The other system has lower sensory precision that generated less accurate responses. The model selects, a priori, the most precise system available to perform the task, therefore, in the absence of any lesions (i.e., pre-morbidly), the model chooses to use the more precise system. Anecdotally, it attends to the task at hand.

We lesioned this model in four different ways and simulated task performance over 1000 trials to compare lesion severity and the time course of recovery. In Lesion 1, we demonstrate that, when the premorbid system is unavailable (after extensively disconnecting extrinsic connections), the alternative system (which is now the most precise) is engaged almost immediately after the synthetic recognises something has changed. The subject is then able to sustain performance with ~60% accuracy after approximately 400 trials. In Lesion 2, we demonstrate the effect of learning in Lesion 1 by showing that the same level of performance is precluded when the capacity to relearn is impaired (by lowering precision in the intrinsic connections). In Lesion 3, we show that incomplete damage to the extrinsic connections in both systems resulted in declining performance with no evidence of recovery—and worse performance than complete damage to one system (i.e., Lesion 1). Finally, the performance of Lesion 3 was even worse when it had reduced capacity to relearn (i.e., Lesion 4). From this we infer that the preserved intrinsic connections in Lesion 3 prevented further decline (i.e., like that observed in Lesion 4).

Our work provides a further step towards integrating the theoretical construct of degeneracy(Sajid *et al.*, 2020c) and clinical patient behavioural data – through in-silico lesions of active inference models, i.e., computational neuropsychology. It offers intuition for what type of functional recovery mechanisms might underwrite particular behavioural profiles, in the context of a word repetition task. This understanding is essential for interpreting changes, in free energy or degeneracy, for word repetition models inverted using patient data(Schwartenbeck and Friston, 2016). Furthermore, the approach is generic and can be applied to other paradigms that investigate language; e.g., picture naming(Sajid *et al.*, 2020a) or speech(Friston *et al.*, 2020).

Future work could investigate the effect of changing the precision of other extrinsic connections. For example, an extrinsic disconnection that renders the Target Word and Spoken Word conditionally independent of each other might show similar performance deficits to those from intrinsic lesion to the Target Word factor. Similarly, recovery via degeneracy could be investigated by disconnecting specific sub-structures of the extrinsic pathways. This would speak to the possibility that biological lesions trigger multiple recovery mechanisms, individually or together, to engender resistance to functional loss. Future studies could also investigate how performance changes when functional reorganisation and re-learning are required to establish a system that was not readily available, (e.g., when the alternative system had extremely imprecise priors before the intervention). In this case, we would expect a sustained drop in performance, imprecise beliefs about the nature of recovery, and longer recovery times, as the model learns to use (i.e., reconfigure via structure learning) an alternative system. Likewise, testing how recovery changes with the degree of asymmetry between the premorbid and alternative models (i.e., imbalanced resource allocation between attentive and inattentive context) may reveal a limited capacity to call upon intact structures and may feature a permanent performance deficit, as observed in chronic patients who fail to fully recover after brain damage.

The impact of other factors beyond lesion type (e.g., age, training intensity) on recovery mechanism (Kleim and Jones, 2008)—would also provide further avenues for interesting future work. For example, the generative model could be equipped with particular priors that enable it to mimic these features—or perhaps it could be given a particular set of experiences (i.e., exposure trials) from which it learns particular parameters. A better understanding of these functional recovery mechanisms—and potential reorganisation processes—may even help target therapeutic strategies after brain insult and improve the effectiveness of rehabilitation (Chua and Kong, 1996; Taub *et al.*, 2002; Berthier and Pulvermuller, 2011).

## 8. Conclusion

Our simulations reveal that recovery depends on the availability of computational and structural resources. While the model could develop resistance to insult, the effects of damage could not be overcome beyond a certain point, leading to persistent impairments. The same model was used to make physiological predictions, by simulating neuronally plausible Bayesian belief updating. The simulated lesions resulted in varied decline in baseline evoked responses. These quantitative predictions indicate the potential of future developments to investigate the neurophysiology of functional recovery and could allow us to infer likely damage based on a patient’s electrophysiological responses and/or functional recovery profile.

## Software note

The generative model in these kind of simulations changes from application to application; however, the belief updates are generic and can be implemented using standard routines (here spm_MDP_VB_X.m). These routines are available as Matlab code in the SPM academic software: http://www.fil.ion.ucl.ac.uk/spm/. The code to replicate this particular generative model and accompanying stimulation data is available: https://github.com/ucbtns/frec

